# Transient BRD4 degradation improves cardiac reprogramming by inhibiting macrophage/Oncostatin M induced JAK/STAT pathway

**DOI:** 10.1101/2023.12.31.573781

**Authors:** Liu Liu, Yijing Guo, Shuo Tian, Ienglam Lei, Wenbin Gao, Zhaokai Li, Shaomeng Wang, Y. Eugene Chen, Zhong Wang

## Abstract

Reprogramming fibroblasts into induced cardiomyocytes (iCMs) holds great potential for cardiac regeneration. However, low conversion rate both in vitro and in vivo remains a significant challenge. To address this challenge, we focused on potential epigenetic barriers and screened 33 small-molecule inhibitors/degraders targeting histone acetyl post-translational modifications and related epigenetic factors. BET degraders were found to enhance cardiac reprogramming efficiency by degrading BRD4 and repressing genes involved in immune response, particularly those in the JAK/STAT pathway. We further identified that macrophage/oncostatin M activated the JAK/STAT pathway to repress cardiac reprogramming. BRD4 degrader treatment improved iCMs formation by inhibiting macrophage/oncostatin M-induced activation of the JAK/STAT pathway. Moreover, BRD4 degrader treatment enhanced MGT-mediated cardiac regeneration in vivo and improved myocardial performance post-myocardial infarction. These findings provide new insights into BRD4, macrophage/oncostatin M and JAK/STAT pathway in fibroblast to cardiomyocyte-like cell conversion and offer promising targets and small molecules to improve iCM reprogramming for clinical applications.

**Highlights:**

1. Screen of histone acetylation modifiers identified BRD4, a histone acetylation reader, as an important regulator in cardiac reprogramming.
2. BRD4 degradation promoted cardiac reprogramming by repressing inflammation and the JAK-STAT pathway.
3. BRD4 degradation inhibited macrophage/oncostatin M-induced activation of the JAK/STAT pathway.
4. BRD4 degradation enhanced MGT-mediated cardiac regeneration in vivo and improved myocardial performance post-myocardial infarction.

## Introduction

Cardiovascular disease (CVD) has remained the leading cause of death worldwide for the past decade^1,2^, with myocardial infarction (MI) being a major contributor in both developed and developing countries. MI leads to impaired heart function due to massive cardiomyocyte loss, fibrosis, and inflammation. While existing clinical treatments for MI have improved patient outcomes, they do not address the underlying issue of cardiomyocyte loss. Direct cardiac reprogramming of fibroblasts into CM-like cells using the Gata4, Mef2C, and Tbx5 transcription factors has emerged as a promising strategy to generate induced cardiomyocytes (iCMs) and replace scar tissue with functional cells^3-5^. However, low conversion rates and limited in vivo application potential pose major challenges for this approach to be used clinically, despite the advantage of utilizing abundant cardiac fibroblasts in the impaired heart.

Cardiac reprogramming involves extensive epigenetic changes, and modulation of these changes has been shown to be an effective strategy to improve cardiac reprogramming efficiency by research from other and our lab^6-9^. However, the role of histone acetylation, as one of the major epigenetic regulations, has not been well-studied in cardiac reprogramming. Histone acetylation is mainly regulated by two groups of enzymes, as histone acetyltransferases (HATs) add, and histone deacetylases (HDACs) remove acetyl groups. In addition, bromodomain-containing proteins specifically bind to acetylated lysine residues and serve as a module deciphering the histone or protein acetylation code.

Our lab has shown that acetylation can be a major target for cardiac regeneration in vivo, as increased histone acetylation induced by VPA and medium chain fatty acid octanoate acid treatments significantly reduces cardiac damage after MI^10,11^. This leads us to question whether we could regulate acetylation to induce cardiac reprogramming and thus reduce cardiac damage after MI in vivo. On the other hand, although several strategies have been applied to address the low conversion rate challenge in vitro, only a few research are focused on mimicking the pathology condition in vivo and test whether the in vitro strategies also applicable in vivo^12,13^. Thus, identifying potential in vivo barriers to cardiac reprogramming and exploring effective strategies, such as using chemical compounds, for in vivo cardiac reprogramming have become research priorities due to their potential for future clinical translation. It would be significant to identify chemical compounds targeting acetylation regulation that have an optimistic effect for in vivo application.

We conducted a screening of histone acetylation-related chemical compound for their effect on cardiac reprogramming. We found that both BRD4 inhibitors and degraders significantly enhanced cardiac reprogramming, resulting in improved conversion of fibroblasts into cardiomyocytes, as measured by cardiac gene expression, sarcomere formation, calcium flux, and spontaneous beating. RNA-seq data revealed significant changes in cell identity-related gene expression, specifically related to immune and fibrosis response pathways, consistent with our previous findings^14^. Notably, a novel BRD4 degrader was found to repress genes involved in immune responses, particularly those in the JAK/STAT pathway. BRD4 degraders protected iCMs from macrophage/oncostatin M (OSM) induced activation of JAK/STAT pathway by affecting BRD4 binding to promoters of those genes. Building on these in vitro results, we applied the BRD4 degrader in a mouse myocardial infarction model, where it effectively induced cardiac-reprogramming-based heart regeneration. These findings suggest that histone acetylation is a promising target for cardiac reprogramming and offers potential applications for translational medicine.

## Results

### 1. Screening of Histone Acetylation-Related Chemicals Identifies BRD4 as a Repressor in Cardiac Reprogramming

To identify potential acetylation epigenetic regulators of induced cardiac myocyte (iCM) reprogramming, we performed a screening of regulators to acetylation regulation. We selected 33 small molecule chemicals related to histone acetylation, including 11 reader inhibitors/degraders, 5 writer inhibitors, 7 eraser inhibitors, and 10 acyl-CoA related metabolites, to target acetylation-related post-translational modifications (Table 1). These chemicals were introduced into an in vitro MGT-induced cardiac reprogramming evaluation system by transducing retroviruses expressing polycistronic Mef2c/Gata4/Tbx5 (MGT) into mouse embryonic fibroblasts (MEFs) derived from a transgenic α-muscle heavy chain (αMHC)-green fluorescent protein (GFP) reporter mouse (Figure 1A). After 14 days of transduction and chemical treatments, the efficiency of reprogramming was evaluated by activation of GFP. We observed 5.49% GFP positive cells in the MGT+DMSO group, indicating a similar success rate of cardiac reprogramming as reported (Figure 1B). From the screening, we are able to identify several chemical inhibitors/degraders that can affect reprogramming efficiency. Among the 33 candidates, several metabolites involved in acyl-CoA metabolism and an inhibitor of HDACs (Trichostatin A), resulted in near-complete abolishment of reprogramming efficiency with certain treatment concentration, while several epigenetic regulators could enhance reprogramming. Among the top 5 chemicals that promoted cardiac reprogramming, 3 were related to acetylation reader inhibition/degradation (Figure 1C). We identified a BET degrader, BETd-246, with the highest efficiency in increasing iCM induction percentage (Figure 1B). Degradation of epigenetic regulators, BET family proteins, by BETd-246 resulted in a 4-fold increase in αMHC-GFP+ cells (Figure 1C), indicating BET family proteins as a barrier to iCM reprogramming.

**Figure 1.**
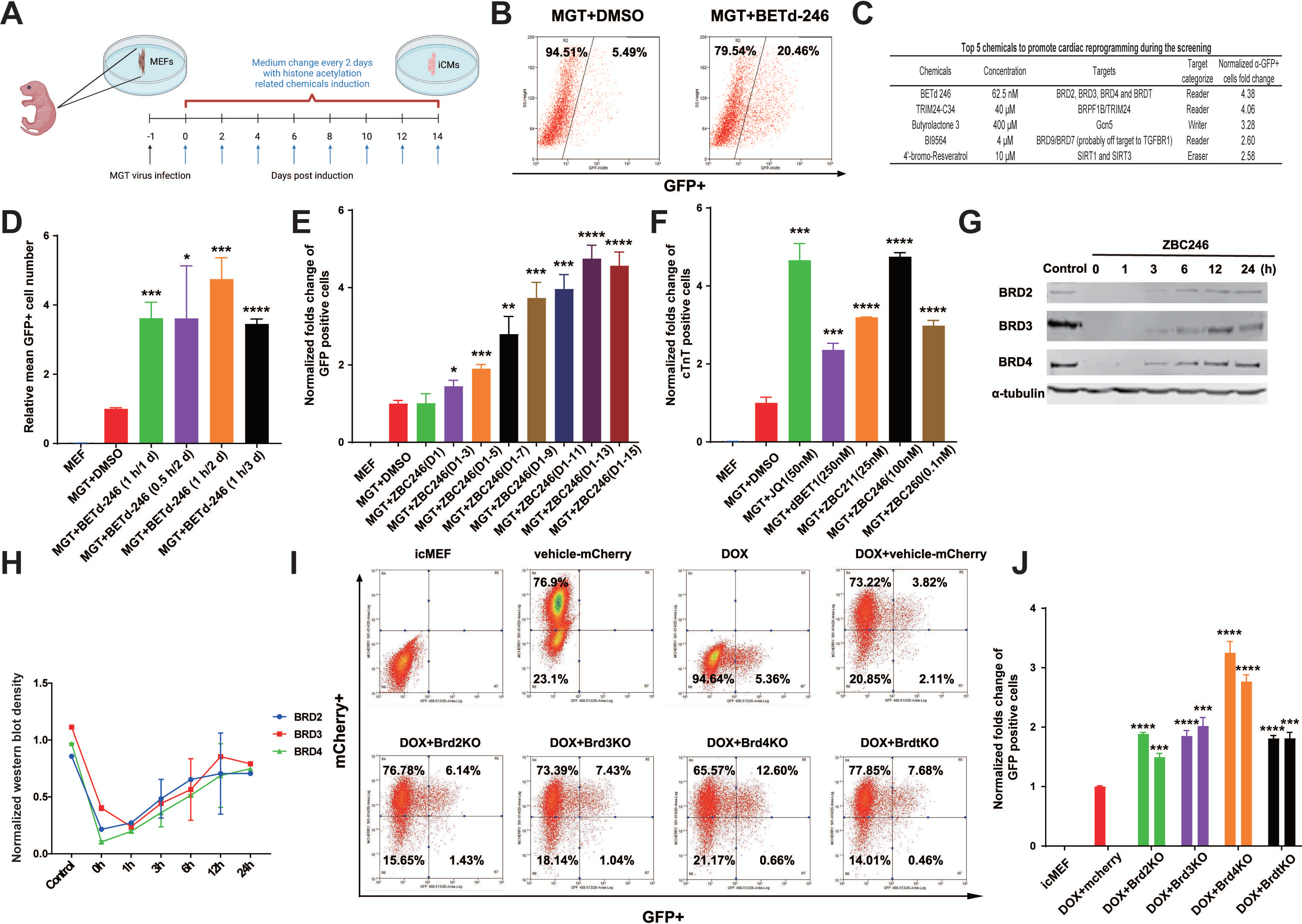
Screening of histone acetylation-related chemicals identified that degradation, inhibition or knockout of BRD4 enhanced cardiac reprogramming. reening of histone acetylation-related chemicals identified that degradation, inhibition or knockout of BRD4 enhanced cardiac reprogramming. A. Schematic illustration of in vitro cardiac reprogramming and histone acetylation related chemicals screening. B. The chemical screening results implied that BET degrader BETd-246 promotes cardiac reprogramming. C. The chemical screening results showed that several modulators promote or inhibit cardiac reprogramming. D. Cardiac reprogramming efficiency under different intervention period and frequency of BETd-246. E. Cardiac reprogramming efficiency under different intervention times of BETd-246. F. Cardiac reprogramming efficiency under different BET inhibitors and degraders intervention. Chemical concentration: JQ1: 50 nM, dBET1: 250 nM, ZBC11: 25 nM, BETd-246: 100 nM, BETd-260: 100 pM. G&H. Protein level of different BET family members under BETd-246 treatment for 1 h. I & J. Cardiac reprogramming efficiency after gene knockout of each BET family members, respectively. mCherry/GFP double positive cells were recognized as iCMs. For each BET protein, two gRNAs for knocking out were used and investigated individually. MGT+DMSO/DOX+DMSO group was used as control group for each experiment. *P < 0.05, **P < 0.01, ***P < 0.001, ****P < 0.0001, compared with MGT+DMSO/DOX+DMSO group. For each group, n = 3.

To further optimize the conditions of BETd-246 in cardiac reprogramming, we tested different treatment durations in our system. Fibroblasts would die after more than continuous 2-hour treatment of BETd-246; therefore, we selected several patterns for BETd-246 treatment and found that 1 hour per two days achieved the highest efficiency (Figure 1D). We also explored whether different treatment times of BETd-246 would affect cardiac reprogramming efficiency. According to our data, the efficiency became higher as the time increased (Figure 1E), indicating the accumulation of reprogramming enhancement by BETd-246. We also compared BETd-246 with several known Brd4 inhibitors and degraders, which showed BETd-246 as the one of the most potent degraders for cardiac reprogramming (Figure 1F).

The degraders used in our research were reported as degraders of BET family members^15,16^. To explore the major effector of BETd-246 in cardiac reprogramming, we first tested the biological function of BETd-246 to its potential targets in our system. Because Brdt is predominately expressed and functions in testicles but not fibroblast^17^, we only tested three other BET family proteins including Brd2, Brd3, and Brd4. Our results indicated that those three proteins were all degraded after 1 hour treatment of BETd-246 in MEFs and gradually increased in the following 24 hours (Figure 1G&H), which means a transient degradation of BET family member can enhance reprogramming. To further determine which BET family member was responsible for the above phenotypes observed in cardiac reprogramming system, each BET family member was knocked out respectively in an inducible MEF cell line called icMEFs line by CRISPR technology. The icMEFs were infected with lentivirus encoding gRNA scaffold, CRISPR-associated protein 9 (Cas9), and mCherry for infection rate calculation. We designed two different gRNA scaffolds for each BET protein and each protein was knocked out in icMEFs, while the reprogramming efficiency in each group was measured by the percentage of mCherry/GFP double positive cell (Figure 1I). We found that the reprogramming efficiency of Brd2, Brd3, and Brdt knock-out group achieved less than 2-fold increase while Brd4 knock-out group achieved nearly 3-fold increase (Figure 1J). This indicates Brd4 as the major BET family member responsible for cardiac reprogramming inhibition and presumably the major effector of the BET degraders.

More importantly, our findings suggest that transient, but not permanent, degradation of BRD4 can enhance reprogramming,since we found that the optimized condition for BET degraders is a 1-hour treatment every two days, but the protein level of BRD4 returns to normal within 24 hours after BETd-246 treatment, those results indicates that a temporary reduction in BRD4 levels is sufficient for enhancing reprogramming.

### 2. Transient BRD4 degradation enhanced cardiac reprogramming of MEFs and NCFs

Next, we further investigated the effect of BETd-246 on cardiac reprogramming of MEFs using the same optimized reprogramming platform. qPCR analysis demonstrated that BETd-246 significantly upregulated the expression of cardiac-related genes (Figure 2A) and downregulated the expression of fibroblast-related genes (Figure 2B). After two weeks of reprogramming, we observed a 5 to 6-fold increase in the expression of cardiac troponin T (cTnT) and α-actinin, as well as a significant increase in the percentage of cTnT/GFP and α-actinin/GFP double-positive cells in the BETd-246 group, as determined by immunofluorescence staining (Figure 2C-F).

**Figure 2.**
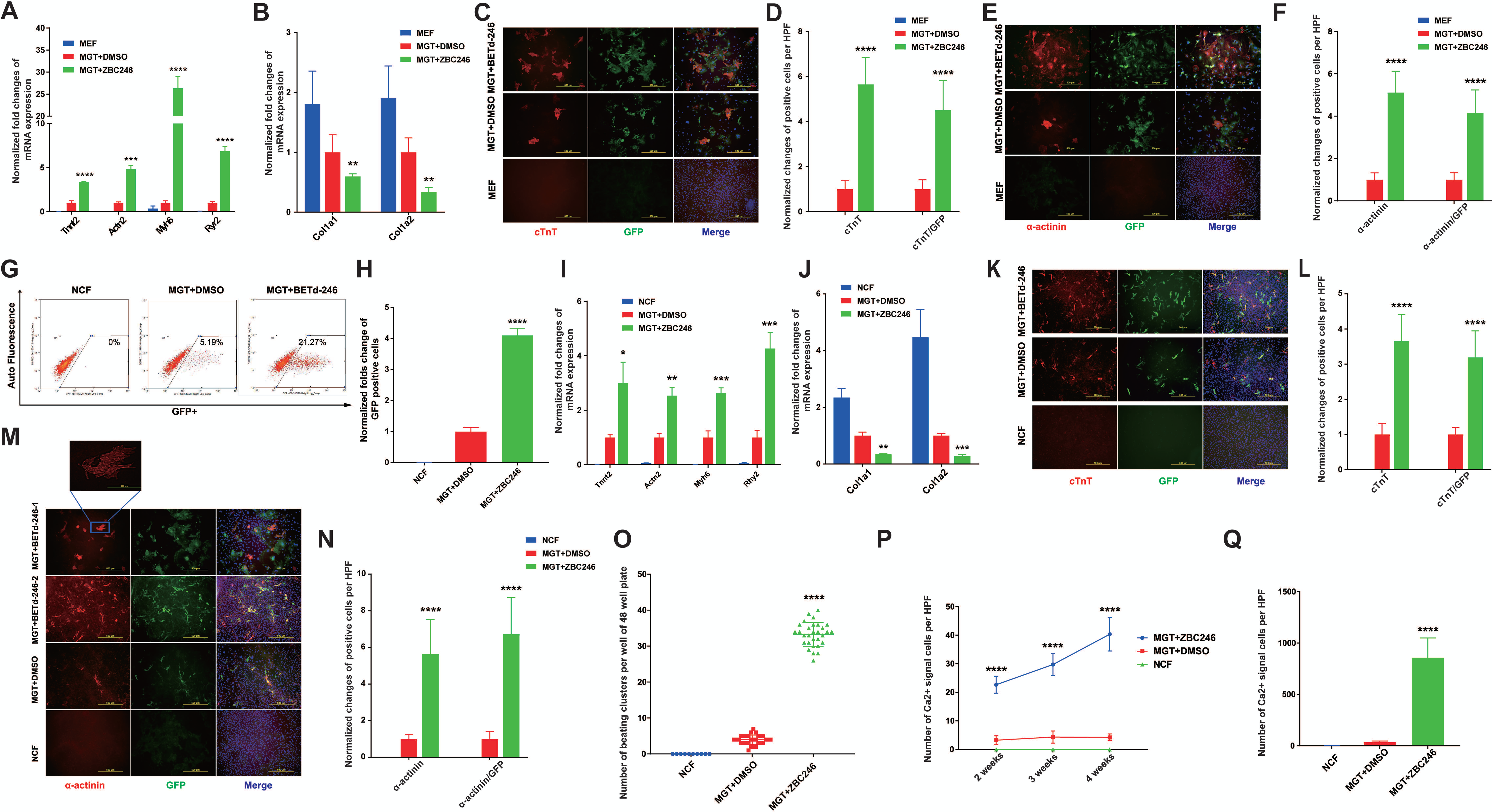
BRD4 degradation enhanced cardiac reprogramming in both MEFs and NCFs. A & B. BETd-246 treatment increased cardiac specific genes expression (A) and decreases fibroblast specific genes expression (B). C & D. BETd-246 treatment increased cTnT+ cell number in MEF-based in vitro cardiac reprogramming. E & F BETd-246 treatment increased α-actinin+ cell number in MEF-based in vitro cardiac reprogramming. G & H. BETd-246 treatment increased cardiac reprogramming efficiency in NCF-based in vitro cardiac reprogramming. I & J. BETd-246 treatment increased cardiac specific genes expression (I) and decreased fibroblast specific genes expression (J) in NCF-based in vitro cardiac reprogramming. K & L. BETd-246 treatment increased cTnT+ cell number in NCF-based in vitro cardiac reprogramming. M & N. BETd-246 treatment increased α-actinin+ cell number in NCF-based in vitro cardiac reprogramming. O. BETd-246 increased the number of beating clusters per well in a 48-well plate in cardiac reprogramming. P. BETd-246 increased the number of cells with intracellular calcium flux in cardiac reprogramming. Q. BETd-246 increased the number of cells with spontaneous calcium transient in cardiac reprogramming. MGT+DMSO/DOX+DMSO group was used as control group for each experiment. *P < 0.05, **P < 0.01, ***P < 0.001, ****P < 0.0001, compares with MGT+DMSO group. For each group, n = 3.

To further evaluate the potential of BETd-246 for cardiac reprogramming, we investigated its effects on neonatal cardiac fibroblasts (NCFs). The MGT+BETd-246 group demonstrated a more than 4-fold increase in induced cardiomyocyte (iCM) formation compared to the MGT+DMSO group, as measured by flow cytometry (Figure 2G&H). Additionally, BETd-246 treatment resulted in increased expression of cardiomyocyte-specific genes (Figure 2I) and decreased expression of fibroblast-specific genes (Figure 2J), as determined by qPCR. Immunofluorescence staining revealed an increase in cTnT and α-actinin expression, as well as cTnT/GFP and α-actinin/GFP double positive cells in NCFs. We could also clearly visualized sarcomere formation in iCMs derived from NCFs treated with BETd-246 but not DMSO (Figure 2K-N).

We further evaluated the function of iCMs, including spontaneous beating and calcium transient. BETd-246-treated NCFs (1 hour per two days) with MGT were cultured for two weeks and then switched to maturation medium for another two weeks. Those cells showed a more than 8-fold increase in iCMs spontaneous beating compared to the MGT+DMSO group (Figure 2O). Spontaneous calcium transient was measured using a genetically encoded calcium indicator and a Rhod-3-staining-based chemically calcium imaging technology. Primary NCFs isolated from αMHC-Cre/Rosa26A-Flox-Stop-Flox-GCaMP3 mice were used for BETd-246-enhanced cardiac reprogramming, which showed a 7-fold increase of intracellular calcium flux of iCMs at two weeks and nearly a 9-fold increase at four weeks compared to the MGT+DMSO group (Figure 2P). Similarly, Rhod-3 staining showed hundreds of cells with spontaneous calcium transient in the MGT+BETd-246 group, while only dozens were observed in the MGT+DMSO group (Figure 2Q).

### 3. Transient BRD4 degradation achieved high reprogramming efficiency of MEFs and NCFs by inhibiting JAK/STAT pathway

We then conducted RNA-seq to analyze changes in gene expression profiles of induced cardiomyocytes (iCMs) after two weeks of reprogramming, with and without treatment with BETd-246 in NCFs and MEFs. Our results showed significant expression changes (p < 0.05 and fold change > 2) in select genes (Figure 3A and Supplement Figure 1A), including those related to cardiac function (Figure 3B and Supplement Figure 1B) and fibroblast markers (Figure 3C and Supplement Figure 1C). In comparison to the MGT+DMSO group, treatment with MGT+BETd-246 resulted in significant downregulation of 284 genes (Figure 3E) and upregulation of 53 genes (Figure 3D)in both NCF-based and MEF-based cardiac reprogramming systems. Gene ontology (GO) analysis of the common regulated genes revealed that BETd-246 treatment stimulated genes related to metabolism, while inhibiting genes related to extracellular matrix organization and immune response (Figure 3D, E). Additionally, GO molecular function analysis suggested that signaling receptor activities were crucial for cardiac reprogramming, while peptidase activity posed a major obstacle (Figure 3D, E). We further found that the JAK/STAT pathway, which has been known to hinder cardiac reprogramming, was downregulated by BETd-246 treatment (Figure 3E). We also identified several JAK/STAT pathways-related genes that were significantly downregulated by BETd-246, prompting us to investigate the link between BET degradation and JAK/STAT inhibition during cardiac reprogramming.

**Figure 3.**
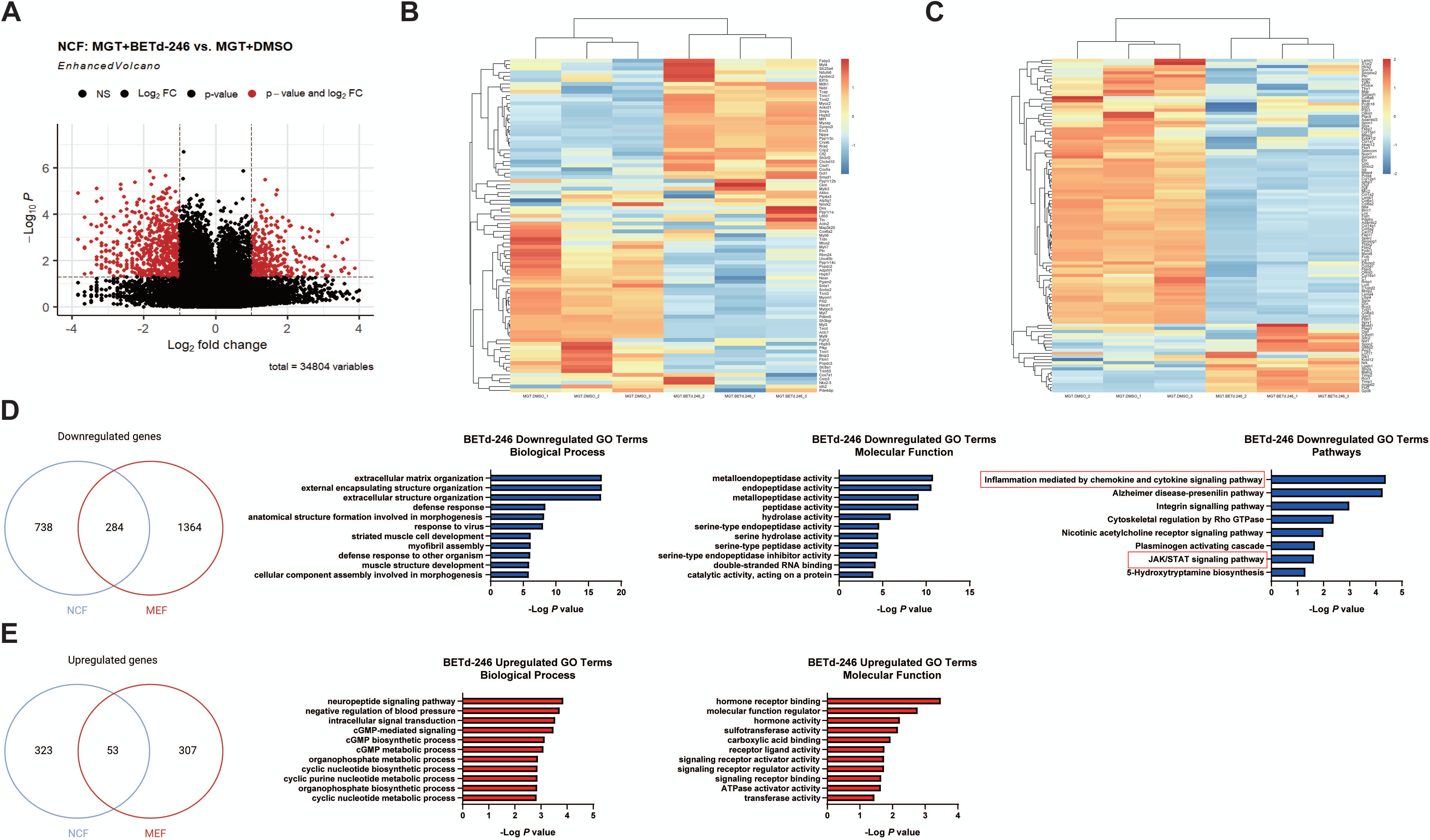
Brd4 degradation achieved high reprogramming efficiency of MEFs and NCFs through inhibiting JAK/STAT pathway. A. Volcano plot showing p-value and fold-change for NCFs treated with MGT+BETd-246 compares with MGT+DMSO. Differentially expressed genes (p-value < 0.05 and FC > 2) are depicted in red. B and C. Heat map showing differential expression of representative cardiac-related (B) and fibroblast-related (C) genes among NCFs. D. MGT+BETd-246 significantly downregulated 284 genes in both NCFs and MEFs (left) compared with the MGT+DMSO group. And top terms from GO biological process analysis, GO molecular function analysis, and GO pathway analysis of those genes were listed (right). E. MGT+BETd-246 significantly upregulated 53 genes in both NCFs and MEFs (left) compares with the MGT+DMSO group. And top terms from GO biological process analysis and GO molecular function analysis of those genes were listed (right). *P < 0.05, **P < 0.01, ***P < 0.001, ****P < 0.0001, compares with MGT+DMSO group. For each group, n = 3.

### 4. BRD4 degradation inhibited macrophage/OSM-activated JAK/STAT pathway to stimulate reprogramming

The activation of JAK/STAT pathway in cells is mostly related to specific transmembrane receptor signals, which are stimulated by extracellular molecules including cytokines. It is acknowledged that inflammation plays an important role in the injury and recovery response of heart after MI, of which JAK/STAT is one of the major target pathways^18^. Macrophage is one of the major sources of cytokines^19^ that involved in acute myocardial injury^20,21^and regulation of repairing process of myocardium during MI. However, whether and how macrophages could affect cardiac reprogramming is less well understood. As results shown above, JAK/STAT pathway was regulated by BETd-246 treatment in our cardiac reprogramming system. Therefore, macrophages may have the capability to regulate cardiac reprogramming via the JAK/STAT pathway. Thus, the effect of macrophages to reprogramming, which is tightly related to the inflammation in heart and activation JAK/STAT pathway during in vivo cardiac reprogramming, was detailly investigated. iCMs were cocultured with macrophages or treated with the supernatant of macrophages after 24 hours incubation, respectively (Figure 4A). Compared with the control medium, macrophages or macrophage supernatant showed significant inhibition to cardiac reprogramming (Figure 4B). Since both macrophages and the supernatant were able to induce reprogramming repression, we believed that paracrine mechanism played an important role in macrophage-induced inhibition to cardiac reprogramming.. Thus, in order to figure out macrophages affect cardiac reprogramming through which cytokines, the major cytokines that were secreted by macrophages, including Angiopoietin-2 (Ang-II), Interleukin-1 beta (IL-1β), IL-3, IL-6, IL-7, Macrophage colony-stimulating factor 1 (M-CSF), Oncostatin-M (OSM), Transforming growth factor beta-1 (TGF-β), and Tumor necrosis factor alpha (TNF-α), was screened systematically in cardiac reprogramming. The screening results showed OSM could completely abolish the reprogramming efficiency (Figure 4C) (Table 2). To further investigate the role of OSM in macrophage-induced inhibition of cardiac reprogramming, an OSM antibody was introduced into the system to neutralize extracellular OSM secreted by macrophages. The results demonstrated that the OSM antibody protected MEF cells from macrophage-induced repression of cardiac reprogramming (Figure 4D). Those results indicated that OSM was the major cytokine that could repress cardiac reprogramming by macrophage.

**Figure 4.**
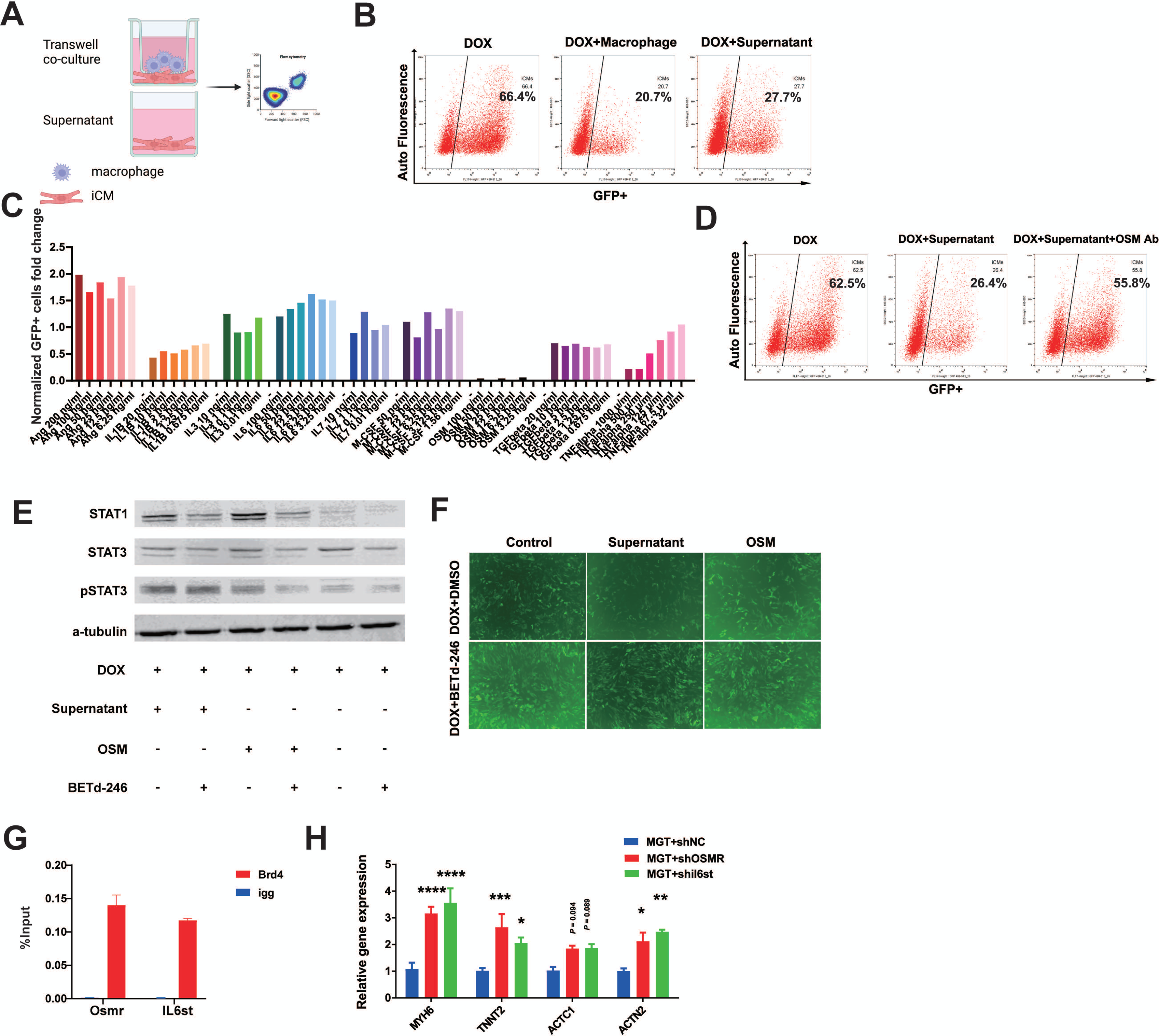
Brd4 degradation inhibited macrophage/OSM-activated JAK/STAT pathway to stimulate reprogramming. A schematic illustration of Macrophage co-culturing and macrophage supernatant treatment. B. Macrophage co-culturing and macrophage supernatant treatment inhibited cardiac reprogramming. C. The cytokine screening results implied that OSM inhibit cardiac reprogramming. D. OSM antibody treatment alleviated macrophage-supernatant-induced inhibition to cardiac reprogramming. E. BET degrader counteracts macrophage supernatant or OSM treatment induced activation of JAK/STAT pathway. F. BET degrader attenuated macrophage supernatant or OSM induced inhibition to cardiac reprogramming. G. BRD4 directly bound to the promoter region of osmr and il6st gene. H. OSMR and IL6ST gene knockdown by shRNA promoted cardiac-specific gene expression in cardiac reprogramming. *P < 0.05, **P < 0.01, ***P < 0.001, ****P < 0.0001, compares with the indicated control group. For each group, n = 3.

Previous studies have shown that OSM can activate JAK/STAT pathway in various cell types^22-24^. In our research, both macrophage supernatant and OSM treatment upregulated the protein level of STAT1, STAT3 and phosphorylation of STAT3 in iCMs (Figure 4E), suggesting that OSM could be a potential JAK/STAT regulator that represses cardiac reprogramming in macrophages. Importantly, under coculture conditions with macrophage supernatant or OSM-treated iCMs, BETd-246 treatment downregulated the protein expression of STAT1 and the phosphorylation of STAT3 (Figure 4E). Furthermore, BETd-246 treatment increased cardiac reprogramming efficiency under coculture with OSM or macrophage supernatant (Figure 4F). These results strongly support the notion that BRD4 mediates macrophage-induced repression of cardiac reprogramming through the JAK/STAT pathway, and its degradation promotes iCM formation from macrophage-induced repression of reprogramming.

Since BRD4 degradation downregulated JAK/STAT pathway gene expression (Figure 3), we hypothesized that BRD4 might directly bind to the JAK/STAT gene regulator regions to control genes expression. To determine how BRD4 regulates OSM-induced signaling events in cardiac reprogramming, chromatin immunoprecipitation (ChIP) assays were performed for several important proteins involved in OSM pathways found by RNA seq analysis. Our results showed that BRD4 was directly bound to the promoters of Osmr and Il6st and affected their gene expression (Figure 4G), which interrupted OSM signaling. OSM receptors including OSMR and IL6ST recognize extracellular OSM and transduce OSM-induced signaling events^25,26^. To further investigate the importance of OSM and its related signaling events in cardiac reprogramming, shRNAs were used to knockdown OSM receptors and interrupt OSM-related pathways. By knocking down of OSMR and IL6ST, respectively, which are the two major OSM receptors, cardiac reprogramming efficiency of MEFs was significantly increased measured by qPCR (Figure 4H). In conclusion, OSM was found to be a key extracellular regulator that inhibited cardiac reprogramming by activating JAK/STAT pathways, while BRD4 counteracted the inhibition from OSM by regulating Osmr and Il6st expression.

### 6. Brd4 degradation enhanced cardiac-reprogramming-based regeneration in vivo

Next, we aimed to investigate whether BRD4 degrader could enhance cardiac regeneration in vivo through cardiac reprogramming. To assess the role of BRD4 degradation in protecting the heart against MI injury, we induced MI in mice by permanent ligation of LAD (Figure 5A). The treatment strategy involved the administration of MGT expression virus to initiate cardiac reprogramming to iCMs, followed by BRD4 degrader treatment initiated 1 week after MI. Cardiac function was evaluated by echocardiography at 4- and 8-weeks post-MI. The results showed that the EF and FS of MI+DSRED group mice were 16.38% and 7.517% respectively, at 4 weeks post-MI while the EF and FS of sham group were 47.31% and 23.55% respectively which indicates a success model induction (Figure 5B and C). While MGT-induced cardiac reprogramming alone slightly improves cardiac function at 4-and 8-weeks post-MI. However, by initiating cardiac reprogramming with MGT expression and improving the reprogramming with BRD4 degrader after MI, significant improvement in EF and FS was observed at 4- and 8-weeks post-MI (Figure 5B and C). BRD4 degradation also lowers infarct size after 8 weeks (Figure 5D and E). These results suggest that continuous administration of BRD4 degrader following MGT-induced cardiac reprogramming can rescue cardiac function after MI (Figure 5F).

**Figure 5.**
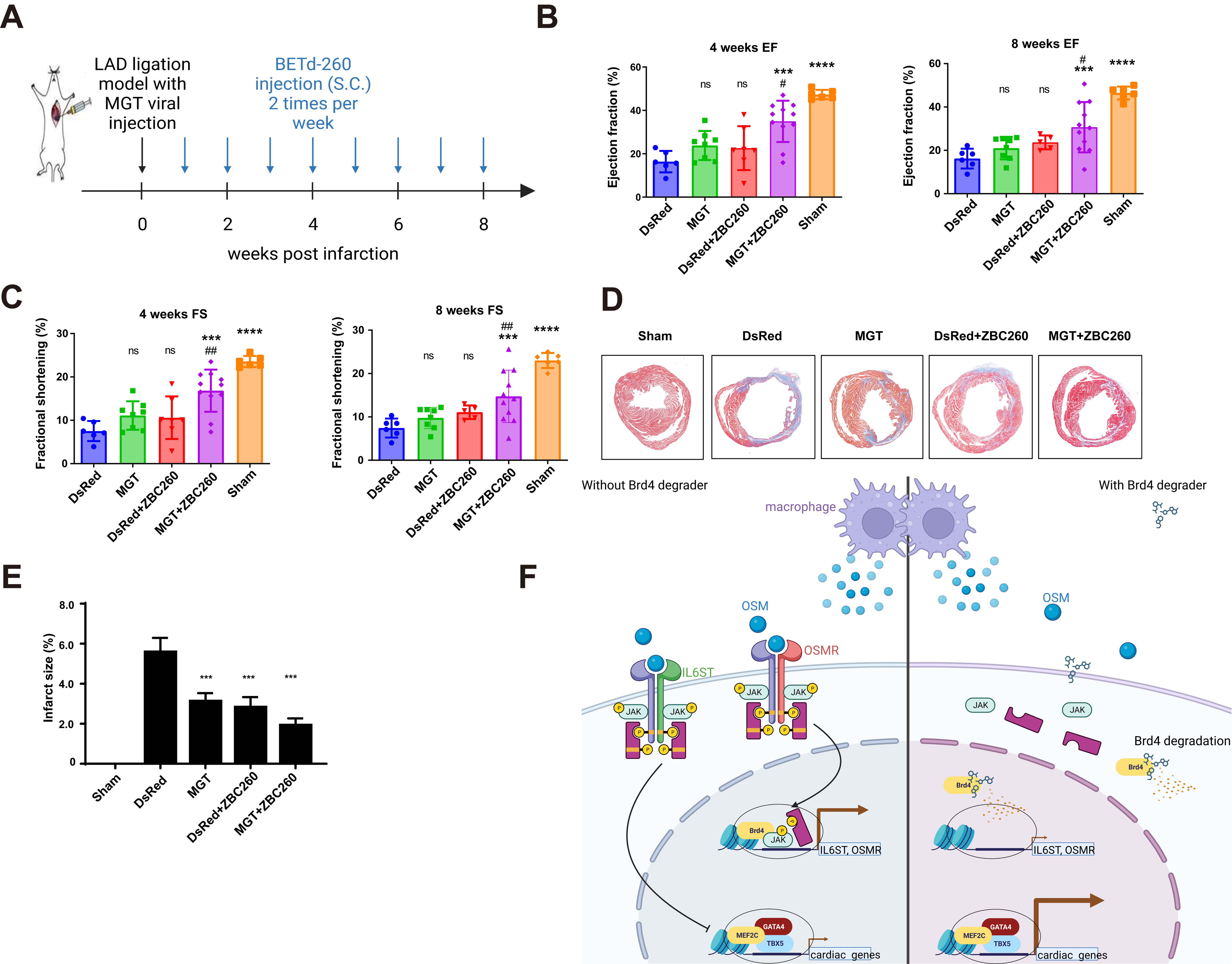
Brd4 degradation enhanced cardiac-reprogramming-based regeneration in vivo. A. A schematic illustration of BET degrader treatment promoted cardiac function in mouse MI model. B. BET degrader improved EF of MI mice 4 (left) and 8 (right) weeks after MI by promoting cardiac reprogramming. C. BET degrader improved FS of MI mice 4 (left) and 8 (right) weeks after MI by promoting cardiac reprogramming. DSRED group was used as the control group. Masson Trichrome staining and calculation of infarct size at 8 weeks (D&E) were performed. (F) A schematic illustration of the mechanism BRD4 degrader attenuates macrophage/OSM-induced reprogramming inhibition. *P < 0.05, **P < 0.01, ***P < 0.001, ****P < 0.0001, compared with DSRED group. For each group, n = 5-11.

## Discussion

In this study, we have shown that the regulation of histone acetylation function through BRD4 degradation enhances cardiac reprogramming both in vitro and in vivo. Specifically, we have identified that a BRD4 degrader BETd-246 improves MGT-mediated cardiac regeneration and improved myocardial performance post-myocardial infarction. Mechanistically, BETd-246 represses macrophage/OSM-induced reprogramming inhibition, leading to downregulation of JAK/STAT pathways. These findings provide new insights into BRD4, macrophage/oncostatin M and JAK/STAT pathway in fibroblast to cardiomyocyte-like cell conversion and offer promising targets and small molecules to improve iCM reprogramming for clinical applications.

Histone acetylation is a crucial regulatory mechanism for cardiac protection, and our study has highlighted the importance of targeting acetylation factors like BRD4 in improving cell fate change. BRD4 has been shown to play a significant role in IPS reprogramming, by removing the signature for the somatic transcriptional program. Our findings suggest that BRD4 is also a key regulator in cardiac reprogramming, and its unique role in cell fate change underscores its importance as a therapeutic target. Our results indicate that the reprogramming process in vitro and vivo is accompanied by the activation of many detrimental pathways, such as JAK/STAT pathways, and BRD4 plays a crucial role in counteracting these pathways, making it a prime target for enhancing cardiac reprogramming.

BRD4 degradation has been studied as a potential treatment for various diseases. ARV-771, a small-molecule pan-BET degrader based on proteolysis-targeting chimera (PROTAC) technology, has been shown to suppress androgen receptor signaling and treat prostate cancer^27^. BETd-246, the degrader used in our in vitro research, has also been proven to be an effective treatment for triple-negative breast cancer^15^. Thus, the safety of using this medication in vivo has already been fully evaluated. Additionally, a previous report has shown that the BRD4 inhibitor JQ1 can have a protective effect on cardiomyocytes after myocardial infarction (MI) by inhibiting inflammation and the TGF-β pathway^28,29^. These studies provide more confidence in the safety of using transient BRD4 degradation in heart regeneration.

One critical distinction to highlight is the differences between in vivo and in vitro reprogramming, particularly the immune response occurred in vivo. Cardiac reprogramming strategies are potential therapies following myocardial infarction, indicating that the entire process occurs within a pathological environment. Macrophages emerge as a pivotal source of cytokines, playing a crucial role in acute myocardial injury and orchestrating the myocardium’s repair during MI. Our findings reveal that macrophages constitute a primary impediment to in vivo reprogramming. Intriguingly, while OSM and IL6 both belong to the IL6 family, we observed no inhibitory effect from IL6, unlike what was noted with OSM. The JAK/STAT pathway has been recognized as a potential inhibitory route, and our results pinpoint macrophages and OSM as upstream activators. Subsequent research will compare the effects of targeting macrophages, OSM, or the JAK/STAT pathway, aiming to optimize the in vivo procedure in conjunction with cardiac reprogramming.

Finally, this study highlights the reversible degradation as a general promising strategy in reprogramming. We believe that a suitable treatment for in vivo cardiac reprogramming therapy should not negatively impact cardiomyocyte viability or function while targeting the major pathway. Ideally, the target protein of such treatment would also have potential cardioprotective effects post-MI. Previous studies have reported that BRD4 inhibition played a protective role in an MI rat model, indicating that BRD4 may meet these requirements as a potential target^30^. These special characteristics of the BRD4 degrader make it an excellent treatment for enhancing the cardiac reprogramming process both in vitro and in vivo. As a result of this special characteristic, this transient degradation of key epigenetic factors likely represents an efficient and safe strategy for in vivo cardiac reprogramming and heart regeneration, with high potential for future clinical applications.

## Methods and Materials

### Mouse lines

The α-MHC-GFP transgenic mice and αMHC-Cre/Rosa26A-Flox-Stop-Flox-GCaMP3 mice were used to derive MEFs and NCFs as described previously^31-33^. All animal related procedures were approved by the Institutional Animal Care and Use Committee of the University of Michigan and are consistent with the National Institutes of Health Guide for Use and Care of Animals.

### Plasmids

The pMXs based retroviral polycistronic vector encoding Mef2c, Gata4, Tbx5 were provided by Dr. Li Qian’s lab^34^. This polycistronic constructs vector was constructed by DNA fragment containing Mef2c, Gata4 and Tbx5 sequentially, which were separated by oligonucleotides encoding P2A and T2A peptides. This polycistronic vector also contains puromycin selection marker for cell purification. Besides, the pMXs based retroviral vector encoding Mef2c, Gata4, Tbx5 respectively were used in animal experiment.

The lentiviral vector named lenti-CRISPRv2-mCherry was acquired from Addgene (99154). This vector contained several DNA fragments including gRNA scaffold, CRISPR-associated protein 9(Cas9) for specific gene knock out and mCherry for infection rate control. psPAX2 (12260) and pMD2.G (12259) was also from Addgene, which were co-transfection with lentiviral vector to generate lentivirus.

### Primary cell isolation

Preparation of MEFs (isolated at E13.5) was previously described^32^. Briefly, embryos of α-MHC-GFP transgenic mice were harvested at 13.5 days post coitum followed by decapitation and removal of internal organs. The tissue was minced and digested with 0.05% trypsin/EDTA (Gibco, Thermo Fisher Scientific). Cells were resuspended in MEFs medium (DMEM medium containing 10% FBS, 1% penicillin/ streptomycin, 10 μl/ml GlutaMAX and 1 mM Sodium Pyruvate) and plated on one 10cm dish per embryo. Then cells were passaged at the ratio of 1:3 (passage 1). Passage 3 MEFs were used for reprogramming.

NCFs were isolated from P2-P3 α-MHC-GFP transgenic or αMHC-Cre/Rosa26A-Flox-Stop-Flox-GCaMP3 mice as reported previously^31,33^. Briefly, heart tissue was isolated, minced and digested with 0.05% Trypsin-EDTA. Then NCFs were collected with type II collagenase (0.5 mg/ml) in HBSS. After washing and resuspending in DPBS with 2.5 g BSA and 0.5M EDTA, cells were incubated with CD90.2 microBeads (miltenyibiotec, Midi MACS Starting Kit) at 4 °C for 30 min. Positive cells were isolated by Magnetic-activated cell sorting (MACS) and plated onto a 10 cm dish with FB medium (IMDM media with 20% FBS and 1% penicillin/ streptomycin) for future use. Isolated fibroblasts were routinely examined under fluorescence microscope or with GFP immunostaining to determine potential CM contamination.

### Inducible fibroblast cell line construction

This inducible fibroblast cell line construction was generated from α-MHC-GFP transgenic mice, called icMEF as reported previously^35^. First, MEFs were isolated from α-MHC-GFP transgenic mice. The primary αMHC-GFP MEFs were therefore transformed with retroviral delivery of SV40 large T antigen and selected transformed cells with Zeocin. Then the Tet-On inducible gene expression system was incorporated into this cell line to permit the temporal regulation of factor expression. The polycistronic MGT construct under the control of a tetracycline responsive promoter for temporal control of MGT reprogramming factor expression was administrated and incorporated into the cell line to generate a new MEF cell line (icMEF) that can be reprogrammed simply by the addition of doxycycline to the culture media.

### Inhibitor information

JQ1 was from Cayman (11187), and dBET1 was from Thermo-Fisher Scientific (NC1087874). ZBC11, BETd-246 and BETd-260 were obtained from Dr. Shaomeng Wang’s lab in University of Michigan.

### Retrovirus and lentivirus preparation

80% confluent 10 cm plates of Plat-E cells were transfected with 10 μg retrovirus vectors by using Lipofectamine 2000 (Thermo Fisher Scientific) plus additional 1.5 ml Opti-MEM (Thermo Fisher Scientific). After 24 hours, the medium was changed with 10 ml fresh MEFs medium. 48 hours and 72 hours after transfection, viral medium was collected twice and filtered through a 0.45-mmcellulose filter. The virus containing medium was added 1/5 vol of 40% PEG8000 solution to make a final concentration of 8% PEG8000. The mixture was kept at 4 °C overnight and spun at 3000 g, 4 °C, 30 minutes to concentrate. The virus was resuspended by fresh MEFs medium with 8 μg/ml polybrene (Sigma).

Similarly, 10 μg lentiviral vectors with 6 μg psPAX2 and 4 μg pMD2.G were packaged into an 80% confluent 10 cm plate of HEK293T cells (ATCC) with 6ml Opti-MEM by using Lipofectamine 2000 (Invitrogen) with another 1.5 ml Opti-MEM (Invitrogen). 4-6 hours later, Opti-MEM was changed by 10 ml fresh MEFs medium. The virus was collected as described before.

### Direct reprogramming of fibroblasts to iCMs

The optimized protocol of direct cardiac reprogramming was described obviously^36^. Briefly, fresh fibroblasts were seeded on tissue culture dishes at a density of 10,000 cells/cm^2^ before virus infection. Fibroblasts were infected with fresh viral mixture containing 8 μg/ml polybrene (Sigma) 24 hours after seeding. Twenty-four hours later, the viral medium was replaced with induction medium composed of DMEM/199 (4:1) (Gibco, Thermo Fisher Scientific) containing 10% FBS, 1% penicillin/ streptomycin and 10 μl/ml GlutaMAX (Gibco, Thermo Fisher Scientific). Medium was changed every 2– 3 days with or without indicated chemicals for two weeks before cells were examined. 1 μg/ml puromycin (SIGP8833-25MG, Sigma) was added into the medium 3 days after infection to eliminate cells without infection.

For spontaneous beating and calcium transient assessment experiments, induction medium was replaced every 2-3 days by mature medium containing StemPro-34 SF medium (Gibco, Thermo Fisher Scientific), GlutaMAX (10 μl/ml, Gibco, Thermo Fisher Scientific), ascorbic acid (50 μg/ml, Sigma), recombinant human VEGF165 (5 ng/ml, R&D Systems), recombinant human FGF10 (25 ng/ml, R&D Systems), and recombinant human FGF basic146 aa (10 ng/ml, R&D Systems) for another 2 weeks as previously described^37^.

### Western blot

Proteins were extracted from cells by adding lysis buffer and centrifuged at 4 °C for 15 minutes at 12,000 rpm. Protein concentration was measured by Bradford protein assay and 40 μg of total protein was separated by SDS-PAGE and then transferred to polyvinylidene difluoride membranes. The membranes were blocked with 5% nonfat dry milk for 1 hour at room temperature and then incubated with primary antibodies over night at 4 °C. After 3 washings with TBST, the membranes were incubated with appropriate secondary antibody in TBST solution for another 1 hour at room temperature. After 3 washings, the membranes were scanned and quantified by Odyssey CLx Imaging System (LI-COR Biosciences, USA).

### Immunocytochemistry

Cells were first fixed with 4% formaldehyde for 15 minutes. Then they were permeabilized with 0.1% Triton X-100 in PBS for another 15 minutes at room temperature. Cells were blocked with 4% horse serum in PBS for 1 hour and then incubated with primary antibodies against cTnT (Thermo Fisher Scientific), α-actinin (Abcam), and GFP (Thermo Fisher Scientific) overnight at 4 °C followed by incubation with appropriate Alexa fluorogenic secondary antibodies (Thermo Fisher Scientific) at room temperature for 1 hours. cTnT, α-actinin, and cTnT/GFP double positive cells were manually quantified by single-blind method from ten randomly HPFs within each well.

### Quantitative real time PCR (qPCR)

Total RNAs from all cells with or without GMT/chemicals induction were extracted using Trizol Reagent (Thermo Fisher Scientific) following the manufacturer’s instructions. RNA integrity was determined using formaldehyde denaturalization agarose gel electrophoresis. RNA concentrations were measured with Nanodrop spectrophotometer (Thermo Fisher Scientific). RNA was reverse transcribed by using iScript cDNA Synthesis Kit (BioRad). qPCR was performed using StepOne Real-Time PCR System (Thermo Fisher Scientific). Primer oligonucleotides were synthesized by Sigma and are listed in Supplementary Table S2.

### Spontaneous beating and calcium transient assessment

Spontaneous beating assessment was performed by light microscopy at room temperature after indicated treatment of NCFs at several time points. Beating cell number was manually quantified by single-blind method after isoproterenol treatment in each well of 48-well plate.

Calcium transient was measured with Rhod-3 Calcium Imaging Kit (Thermo Fisher Scientific) according to the manufacturer’s instructions. αMHC-Cre/Rosa26A-Flox-Stop-Flox-Gcamp3 NCFs were also used for calcium transient assessment as previously reported^38^ that was performed by fluorescence microscopy at room temperature after indicated reprogramming treatments. Three HPFs of view were randomly selected within each well, video was recorded for 3 minutes and manually quantified by single-blind method.

### RNA-sequencing

Total RNAs from all groups of cells were isolated using TRIzol following the provider’s instructions. RNA (RIN > 8.5) was used for RNA-seq library preparation by using NEBNext® Ultra™ II Directional RNA Library Prep Kit for Illumina (E7760S). The libraries were sequenced using Hiseq 4000 by the University of Michigan Sequencing Core. The quantification of RNA expression was estimated by Kallisto ^39^. Differential gene expression analysis was done using the R package DESeq2. The abundance of genes was used to calculate fold change and p values. Cutoff values of fold change greater than 2 and p values less than 0.05 were then used to select for differentially expressed genes between sample group comparisons.

Significant pathway enrichment analysis was performed using PANTHER Overrepresentation Test (release 20150430, http://geneontology.org)^40,41^.

### Animal experiment

All experiments were approved by the Animal Care and Use Committee of the University of Michigan and were performed in accordance with the recommendations of the American Association for the Accreditation of Laboratory Animal Care.

Myocardial infarction (MI) surgeries with or without virus injection were performed as previously reported with minor optimization^10^. Briefly, mice were anaesthetized with ketamine (100 mg/kg) and xylazine (10 mg/kg). Myocardial ischemia was performed by permanent occlusion of the left descending coronary artery (LAD) using 8-0 silk sutures. For MI studies, mice were randomly divided into 5 groups: Sham, MI+DSRED, MI+DSRED+BETd-260, MI+MGT, MI+MGT+BETd-260. The retrovirus encoding MEF2C, GATA4, TBX5 or DSRED (control) respectively was packaged in Plat-E cells as we described above. Retrovirus was collected 48 and 72 hours after transfection and purified by sucrose density gradient ultracentrifugation. The titer of retrovirus was determined by qPCR. Except for Sham group, retrovirus was directly injected into the infarcted area of heart following LAD occlusion during the surgery by 10^8^ pfu/mouse. Mice in MI+DSRED+BETd-260 group and MI+MGT+BETd-260 group were injected by subcutaneous with BETd-260 (5mg/kg) 2 times per week for 4 weeks. The mice in other groups were administered with a corresponding dose of saline. At the indicated time points, mice were euthanized for assay. Additionally, it was found that initiating BRD4 degrader in the acute phase did not improve heart function (data not shown).

Echocardiography was performed before and after surgery at the indicated time points. Left ventricular internal diameter end diastole (LVIDd) and end systole (LVIDs) were measured perpendicularly to the long axis of the ventricle. Ejection fraction (EF) and fractional shortening (FS) were calculated according to LVIDd and LVIDs. All echocardiography measurements were performed by a single-blinded investigator. Histological studies were performed as previously described^10^. Briefly, mice were sacrificed, and the hearts were perfused with 20% KCl. After fixed with zinc fixative solution (BD Pharmingen) and dehydrated by alcohol, the hearts were embedded by paraffin and sectioned into 5 μm slides. The sections were processed for Masson’s trichrome. Images were captured by Aperio (Leica Biosystems, Buffalo Grove, IL, USA). After Masson’s trichrome staining, the epicardial infarct ratio was measured by dividing epicardial infarct lengths by the normal epicardial length of left ventricle. Endocardial infarct ratio was similarly measured. Infarct size was calculated as [(epicardial infarct ratio + endocardial infarct ratio) /2] × 100%^42^.

The LV tissue was collected and total RNAs were extracted by Trizol following manufacture’s protocol. qPCR was then performed as described above.

### Statistical analysis

Results were presented as mean ±s.e.m. Statistical difference between groups was analyzed by one-way ANOVA followed by the Student–Newman–Keuls multiple comparisons tests. A P-value <0.05 was regarded as significant. Each experiment was performed at least twice.

---

Supplement Figure 1. RNA-seq data in MEFs. A. Volcano plot showing p-value and fold-change for MEFs treated with MGT+BETd-246 compares with MGT+DMSO. Differentially expressed genes (p-value < 0.05 and FC > 2) are depicted in red. B and C. Heat map showing differential expression of representative cardiac-related (B) and fibroblast-related (C) genes among MEFs.

